# Varieties of the plant *Ctenante oppenheimiana*: A colorful enigma of evolution

**DOI:** 10.1101/300376

**Authors:** Humberto A. Filho, Odemir M. Bruno

## Abstract

The plant *Ctenante oppenheimiana* presents an interesting contrast of colors in the abaxial epidermis. A striking feature is that the stomata are green and cover a purple pavement where the pavement cells have green cellular walls. These characteristics have been used in studies about ecophysiology and phenotypic plasticity. However, the existence of a second tricolor variety of the plant make these characteristics even more heterogeneous and introduce new paradigms of the physiological role underlying this morphology. In this work we show by optical microscopy images the striking differences between the varieties bicolor and tricolor of the plant *Ctenante oppenheimiana*

## Introduction

The Ctenante genus was described in 1883 [7], almost a century later the plant was classified in the marantacea family, [3]. The 16 known species of Ctenanthe are distributed mainly along the brazilian atlantic coast, with a few representatives in the Amazon region of Brazil. Ctenanthe oppenheimiana, is a species of flowering plant with adaxial and abaxial surfaces of the leaves green and purple respectively. Interestingly, exist a variety tricolor from this specie that presents a color morphism, on both the abaxial and adaxial surfaces, where whiter shades of purple and green composes the surface, forming a mosaic of colors on each side. [12].

A purple coloration of lower abaxial leaf surfaces is commonly observed in deeply-shaded understorey plants, especially in the tropics. However, the physiological role of the pigmentation, including photosynthetic adaptation, remains unclear [10]. The pigments responsible for the coloring are known by anthocyanins, it can be found in plant species across a broad range of habitats, especially common in understorey plants of the tropics.

Although the distribution of abaxial coloration among tropical taxa are widespread, it is not known the ecophysiological aspects of this phenotype. The photoprotection have been implicated in the rules played by anthocyanins in plants, in which exposed abaxial leaf surfaces are vulnerable to high-incident light. [9] Some studies have observed that the purple abaxial leaf surfaces appeared to reflect more red light than green surfaces. [11]. This observation induces the hypothesis that anthocyanin pigments may reflect adaxially-transmitted red light back into the mesophyll, in order to capture red photons in environments where light is limiting. This hypothesis is known as back-scatter propagation, it has not yet received a complete experimental validation but could explain the color of leaves in understorey plants. It is known that the abaxial surface of the leaves, in some species of purple plants, presents a high concentration of green stomata cells, which generates an interesting contrast between the color of the pavement of the epidermis (purple) and the stomata (green), making this cells completely visible in studies of density and distribution pattern on the leave epidermis. [6]. The stomatic density varies according to the environmental light exposure. [8], then the color of the epidermis may also influence the distribution of stomata. The striking green color of both the stomata and the cell walls of the pavement cells, contrasting with the purple color of the rest of the epidermis floor, raises questions about the ecophysiological and evolutionary relationships of the presence of chlorophyll in such a high amount in these structures [2] In this work, we discussed the incredible color variation of the abaxial epidermis and leaf pavement in two varieties of *Ctenante oppenheimiana* and the color conservation of cell walls and green stomata in these varieties, through optic microscopy images of the abaxial epidermis. We are looking for hypotheses for the maintenance of tricolor variety in the nature and the possible evolutionary advantages of the variability of colors in the leaf. The phenomena of light capture in different frequencies, heat reflection, photosynthesis in the cell wall at leaves with different shades of purple and green, between these varieties, can throw new paradigms to the study of photosynthesis.

## Materials and Methods

### Plant material

The *Ctenante oppenheimiana* varieties with tricolor and bicolor leaves, were analyzed by optical microscopy. Samples of 0.5 *cm*^2^ of the leaves in each plant were put in slides, with the abaxial surface facing up, covered with water, over covered with coverslips, and submitted to optic microscopy at microscope Axio-Lab A1-Zeiss at magnifications 50× and 100×.

Microscopic and whole plant images, from tricolor and bicolor varieties are available respectively at the following web addresses:

https://dataverse.harvard.edu/dataset.xhtml?persistentId=doi:10.7910/DVN/DNVSO1 [5]

https://dataverse.harvard.edu/dataset.xhtml?persistentId=doi:10.7910/DVN/EN6BBG [4]

## Results

The abaxial surface of the two-color variety of *Ctenante oppenheimiana* showed marked characteristics as to the color of the stomata (green) relative to the color of the purple pavement. The color contrast is incredible. Optical microscopy also makes evident the green coloration of the cellular walls of the pavement cells (see figure 2). These characteristics make this plant a model for studying stomata and distribution of stomata associated with environmental variations and raise questions about the physiological functions related to observed morphology. It is also evident that the *Ctenante oppenheimiana* plant can be used as a model plant in morphometric studies based on distances between stomata [1]. The figure 2 shows an optical microscopy image from the abaxial epidermis of the plant and a model to measurement of the phenotipyc plasticity based on distance between the stomata from abaxial surface. 2.

**Figure 1.**
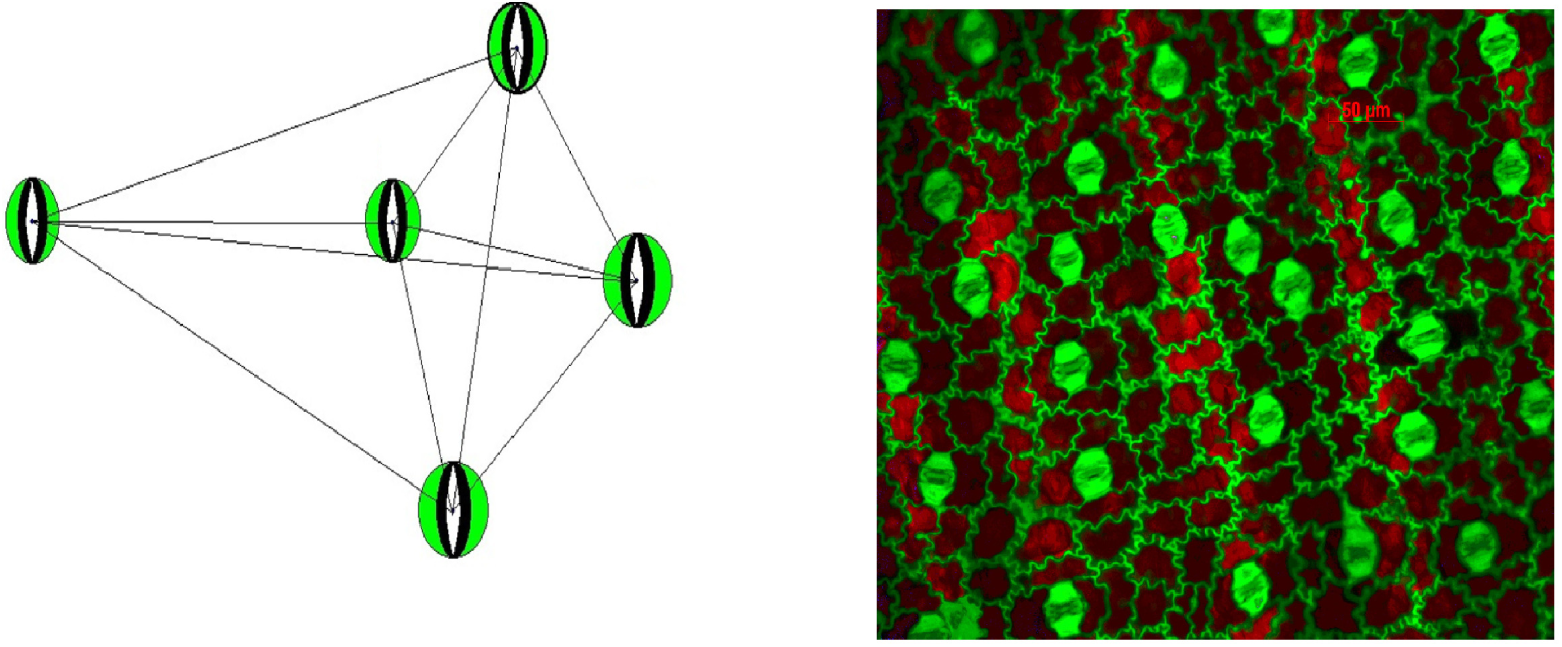
A) Connection network of the stomata neighbours, the edges represent the distance between stomata B) Optic microscopy from abaxial leaf from *Ctenante oppenheimiana*, magnification 200×, highly contrasting green stomata and pavement cell wall.

**Figure 2.**
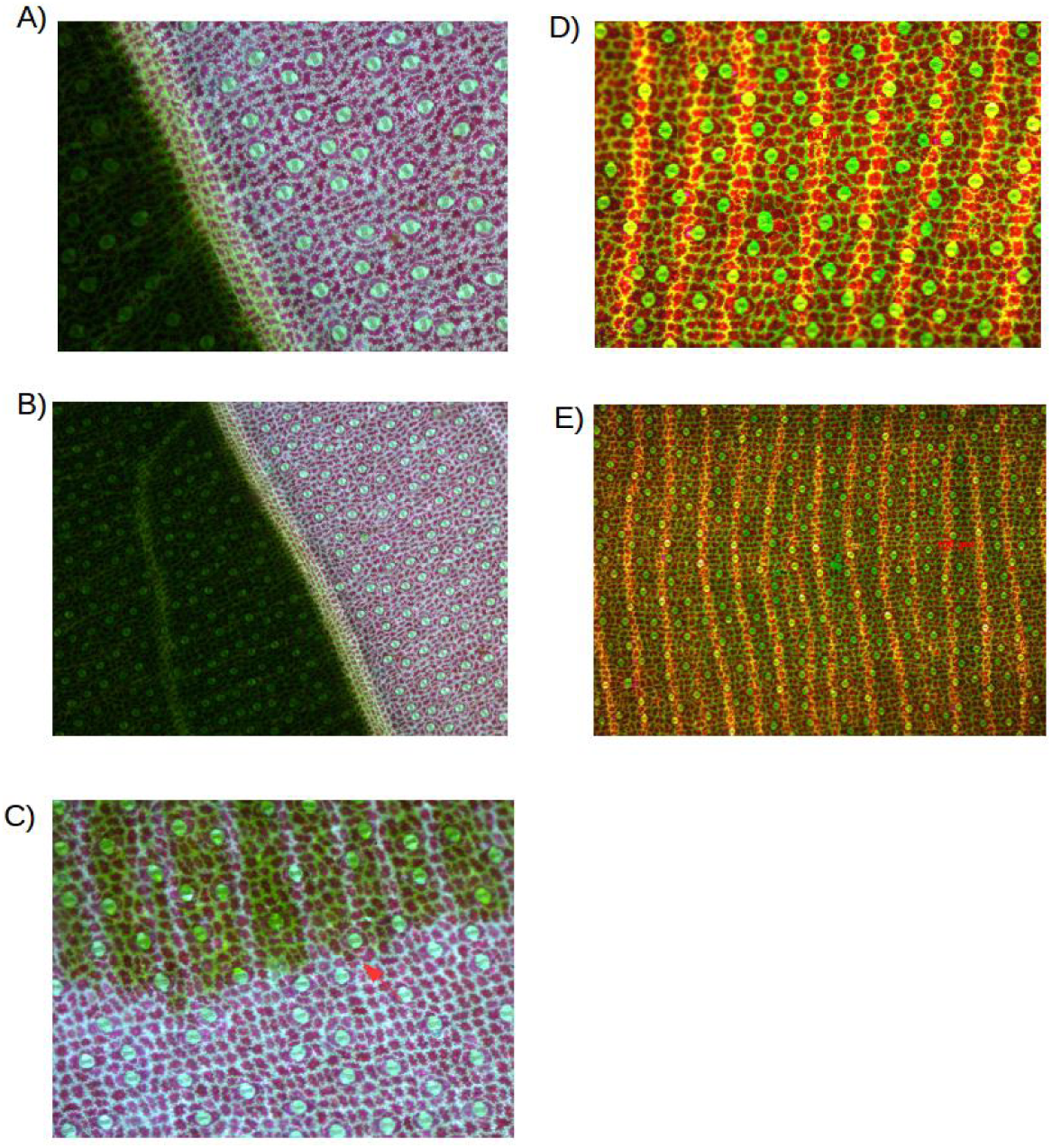
A) Abaxial surface from tricolor variety magnified 100×, transition areas with high and low chlorophyll content at superior epidermis. Green stomata and purple pavement cells with green cellular wall. B) Abaxial surface from tricolor variety magnified 50× C) Transition area at leaf from tricolor variety with high and low chlorophyll distribution at superior epidermis. The red arrow indicates the transition (magnification 100×). D) Abaxial leaf from bicolor *Ctenante oppenheimiana*, magnified 100×, highly contrasting green stomata and pavement cell wall. There is no transition areas with evident differences in chlorophyll at superior epidermis E) Abaxial leaf from bicolor *Ctenante oppenheimiana*, magnified 50×

The existence of the tricolor variety of the *Ctenante oppenheimiana* raises the issue of morphological differences and heterogeneities between varieties. Images obtained with optical microscopy images of the abaxial surface of the tricolor variety, shows that the amount of chlorophyll pigment in the upper layer of the abaxial epidermis, varies according with different leaf area. In some cases the high amount of chlorophyll covers the purple surface causing a dark greenish that recovers the color purple (see figure 2.

## Discussion

The comparison of the microscopic morphology between *Ctenante oppenheimiana* varieties open questions that require studies on photosynthesis, habitat ecology and adaptation, genetic markers of color, phenotypic plasticity and physiology. At the same time the observation of such contrasting characteristics in the abaxial epidermis of this species makes it a model for the study of all these subjects.

## Acknowledgments

H.A.F. gratefully acknowledges the financial support of CNPq grant 153137/2013-4.

